# Towards Similarity-based Differential Diagnostics For Common Diseases

**DOI:** 10.1101/2021.01.26.428269

**Authors:** Luke T Slater, Andreas Karwath, John A Williams, Sophie Russell, Silver Makepeace, Alexander Carberry, Robert Hoehndorf, Georgios V Gkoutos

## Abstract

Ontology-based phenotype profiles have been utilised for the purpose of differential diagnosis of rare genetic diseases, and for decision support in specific disease domains. Particularly, semantic similarity facilitates diagnostic hypothesis generation through comparison with disease phenotype profiles. However, the approach has not been applied for differential diagnosis of common diseases, or generalised clinical diagnostics from uncurated text-derived phenotypes. In this work, we describe the development of an approach for deriving patient phenotype profiles from clinical narrative text, and apply this to text associated with MIMIC-III patient visits. We then explore the use of semantic similarity with those text-derived phenotypes to classify primary patient diagnosis, comparing the use of patient-patient similarity and patient-disease similarity using phenotype-disease profiles previously mined from literature. We also consider a combined approach, in which literature-derived phenotypes are extended with the content of text-derived phenotypes we mined from 500 patients. The results reveal a powerful approach, showing that in one setting, uncurated text phenotypes can be used for differential diagnosis of common diseases, making use of information both inside and outside the setting. While the methods themselves should be explored for further optimisation, they could be be applied to a variety of clinical tasks, such as differential diagnosis, cohort discovery, document and text classification, and outcome prediction.

## Introduction

While a great effort has been invested in the digitisation of healthcare information and data, natural language remains the primary mode of communication and record in the domain. As such, electronic health records (EHRs) contain a wealth of natural language text that comprises an important and fruitful resource for secondary uses, leading to important insights and improved outcomes for patients [1, 2].

Biomedical ontologies encode knowledge in the healthcare domain, including phenotypes and diseases. For example, the Human Phenotype Ontology (HPO) describes disease phenotypes observed in humans, and is used to encode phenotype profiles for real-world entities in healthcare, including patients and diseases [3]. Phenotype ontologies can then be employed to describe phenotype profiles for biomedical entities by associating sets of ontology classes with those entities. Biomedical ontologies have semantic features, including a taxonomic classification, which can be represented as a directed acyclic graph, in which directed edges encode specification of concepts by transitive subsumption relationships.

These resources have been widely explored for the purposes of differential diagnostics and identification of candidate genes underlying rare diseases. In particular, measurements of semantic similarity between phenotype profiles have been used to generate hypotheses and predict diagnoses [4]. Semantic similarity is a measure of how similar two classes are, derived from analysis of the taxonomic structure of an ontology [5]. Measures of similarity can also be determined for groups of classes, including phenotype profiles. The intuition is that a patient phenotype profile will be more similar to the phenotype profile describing the disease they actually have, than to those they do not. This expectation can be leveraged to produce diagnostic hypotheses.

The Online Mendelian Inheritance in Man (OMIM) and OrphaNet databases contain phenotype profiles for many rare genetic diseases [6, 7], manually curated from literature. Semantic similarity has been used to successfully classify rare disease patients by comparing their phenotype profiles, produced either by clinical experts or patients themselves, with those in OMIM and Orphanet [8]. These approaches form the basis of a useful differential diagnostic and clinical decision support methodology [4].

Phenotype similarity methods have been extended to involve data derived from clinical text narratives [9]. Doc2HPO [10] explored the creation of patient phenotypes using HPO from clinical narratives, using text mining to aid experts in curating phenotype profiles which could then be used directly in various differential diagnosis and gene prioritisation tools. Another work developed an approach for deep phenotyping patients from clinical narratives, producing HPO phenotype profiles, particularly for the purpose of disease gene variant prioritisation [11]. Other work has focused on expanding and enriching the phenotype profiles described by OMIM and OrphaNet using EHRs [12]. Finally, literature mining of relationships between DO and HP has been employed to identify relationships between diseases and phenotypes [13], again for variant prioritisation.

Semantic similarity has also been used to support diagnosis and clinical decisions within particular disease domains, such as in skeletal dysplasia [14] and breast pathology [15]. These approaches rely on the use of extensive and specific application ontologies, rather than general phenotype ontologies.

Aside from similarity-based methods, machine learning with quantitative features, as well as features derived from text directly or in the form of word embeddings, have been employed for rare disease diagnosis over EHRs [9, 16]. Differential diagnosis of diseases through conversion of clinical pathway definitions to sets of rules mapped to ontologies has also been previously been explored, but did not involve clinical narratives or semantic similarity [17].

While semantic similarity has proven a successful approach for classification of rare diseases through comparison with sets of phenotype profiles, it has not been applied to common diseases or generalised diagnostics in a clinical setting. Part of the reason for this is a lack of definitional phenotype profiles for common diseases. Definitional phenotype profiles are a list of phenotypes associated with a disease, which are typical for patients with that disease to have. For example, OMIM provides definitional phenotype profiles for rare diseases, describing phenotypes typically observed in those patients, and which indicate that patient may have that disease.

We developed an approach for generalised common disease diagnosis, exploring solutions to the problem of limited availability of disease–phenotype associations. First, we create phenotype profiles for patient visits using concept recognition over text associated with patient visits from MIMIC-III. We then evaluate and contrast two methods of predicting diagnosis: comparison to other patients in the cohort, and secondary use of definitional phenotype profiles previously mined from literature. We also consider a combined method, by selecting and annotating a training set of 500 patient visits, whose text-derived patient phenotypes we use to extend the literature-derived phenotypes. In doing so, we evaluate whether similarity-based approaches are feasible for general use in common disease classification, whether uncurated text-derived phenotypes can be used for semantic similarity classification, whether text-mined phenotype profiles for diseases from literature are applicable in a clinical setting, and whether those profiles can be improved by in-domain training.

## Methods

### Data Preparation and Information Extraction

MIMIC is a freely available healthcare dataset, describing nearly 60,000 critical care visits across three hospitals with a combination of structured and unstructured data, including textual notes [18]. Within MIMIC, diagnoses are provided in the form a canonical ICD-9 code, produced in the original care setting by clinical coding experts. We sampled 1,000 patient visits from the MIMIC-III dataset, collecting their associated texts together into one file per patient visit. We limited our patient visit sample to those with a primary ICD-9 diagnosis that mapped to a class in the Disease Ontology (DO) [19], since the definitional disease phenotypes we used are associated with DO classes. Mappings were obtained from DO, using its in-built annotation properties that define database cross-references.

We then used the Komenti semantic text mining framework [20] to create a vocabulary from all non-obsolete terms in HPO. Subsequently, we used Komenti to annotate the texts associated with each sampled patient visit, producing in effect, a list of HPO terms associated with each patient visit, or a phenotype profile for each patient visit.

To obtain a set of definitional disease phenotypes, we re-used a dataset from a previous experiment which derived phenotypes for DO diseases by text-mining literature abstracts for class co-occurrence [13]. We pre-processed this dataset to remove all non-HP phenotypes from the definitions, and collapsed multiple definitions into single definitions with all unique HP classes.

To create a training set for in-domain optimisation, we also sampled 500 patients who were not included in the set of 1,000 patients used for evaluation. Using the same annotation method, we also produced phenotype profiles for these patient visits. To create a separate set of optimised disease profiles, we extended the literature disease profiles with all unique HP associations found for patient visits with matching primary diagnoses in the training set.

### Semantic Similarity and Evaluation

We then produced, for each patient visit, three sets of similarity scores. The first set of similarity scores is formed of pairwise similarity with each other patient visit phenotype profile in the sample. The other two were formed of pairwise similarity scores between patient visit phenotypes and the literature-derived disease phenotypes, with and without the in-domain optimisation from the training set. The difference between patient-patient comparisons and patient-disease comparisons is demonstrated in Figure 1.

**Figure 1.**
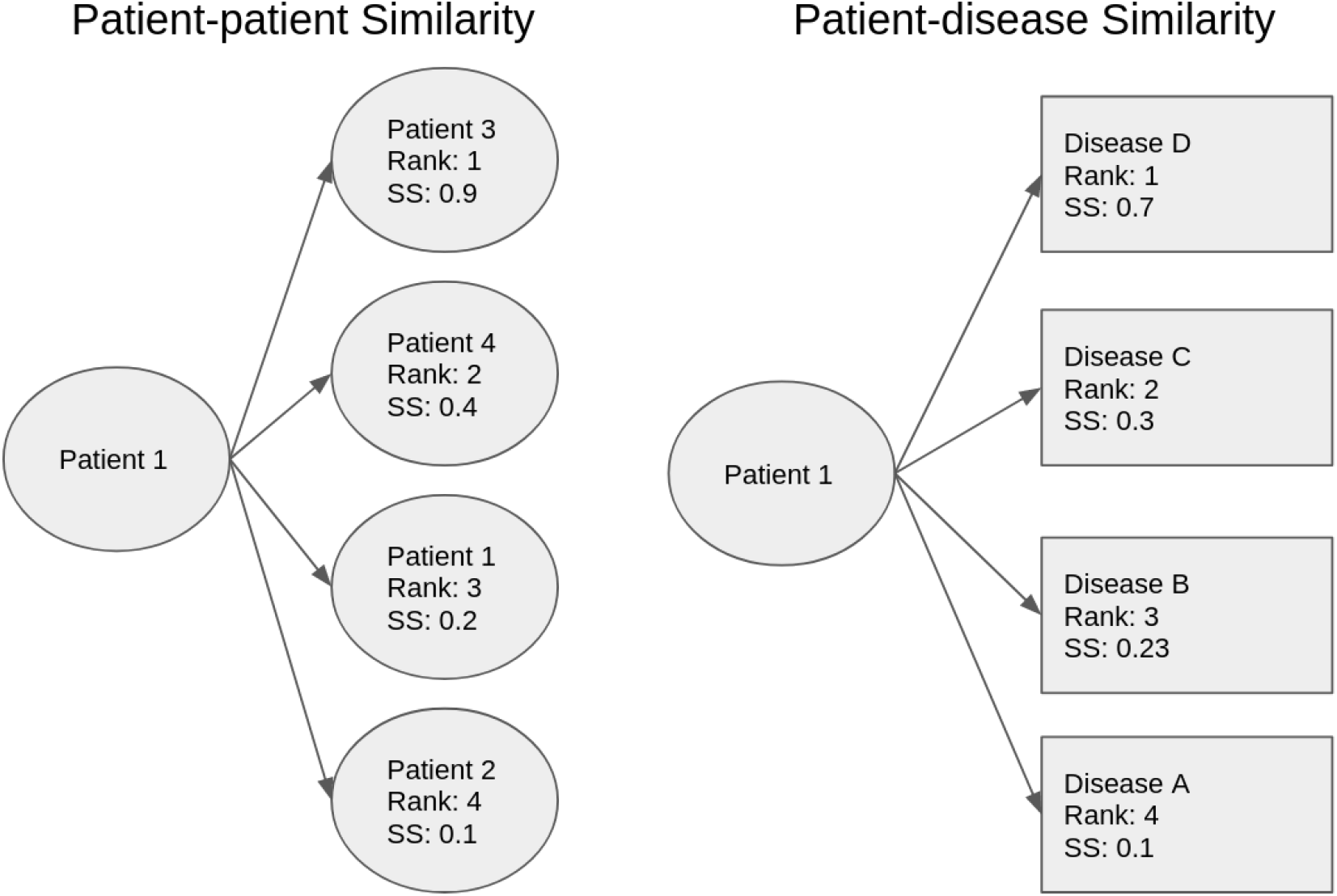
Comparison between the two methods for generating rankings for each patient. Each box signifies a phenotype profile comparison based on an example where there are five patients and four disease profiles. On the left side, the patient’s profile is compared with each other patient profiles in the set, and the ranking is produced from the semantic similarity score between a patient’s profile and every other patient profile. On the right side, the patient’s profile is compared with every disease phenotype profile, and the ranking is produced by ordering the semantic similarity score between the patient and every disease profiles.

To calculate semantic similarity scores, we used the Semantic Measures Library [21]. We used the Resnik measure of pairwise similarity and information content calculated from annotation frequency [22], with Best Match Average for groupwise similarity [23]. In the case of the patient-patient similarity calculations, information content was calculated using a corpus of all HPO codes associated with sampled patient visits. For the purpose of patient visit-disease phenotype similarity calculations, information content was calculated using all HPO codes associated with sampled patient visits, and all HPO codes associated with all disease phenotypes.

We then measured the ability of ranked similarity scores to be predictive of primary diagnosis. We evaluated the results using Area Under the receiver operating characteristic Curve (AUC), Mean Reciprocal Rank (MRR), and Top Ten Accuracy (the percentage of patients for whom the correct diagnosis was in the top ten most similar entities). The software for the experiment is freely available at https://github.com/reality/miesim.

## Results and Discussion

We created phenotype profiles for 1,000 patient visits sampled from MIMIC-III by associating them with HPO terms identified in their text narrative. We then explored two different methods of classifying primary patient diagnosis: comparison with other patients, and comparison with literature-derived disease phenotype profiles (described in Figure 1). To evaluate whether these methods could be combined, to lead to improved performance, we also created an extended set of disease phenotypes, by extending the literature-derived phenotypes with HPO terms mined from the clinical narrative text associated with a training set of 500 patient visits.

Table 1 summarises the results for use of semantic similarity to predict primary diagnosis using patient-patient comparisons and patient-disease profile comparisons, while Figure 2 compares the ROC curves. The AUC metric shows that, of the patient comparison and disease comparison settings, comparison with disease phenotypes produces a better overall classifier.

**Table 1.** Performance for matching first diagnosis of MIMIC patients under different settings. Top ten accuracy is the percentage of patients for whom a correct diagnosis appeared in the ten most similar entities.

| Setting | AUC | MRR | Top Ten Accuracy |
| --- | --- | --- | --- |
| Patient Comparison | 0.774 (0.7724-0.7762) | 0.423 | 0.606 |
| Disease Comparison | 0.869 (0.8564-0.8818) | 0.016 | 0.029 |
| Trained Disease Comparison | 0.974 (0.9682-0.9796) | 0.314 | 0.638 |

**Figure 2.**
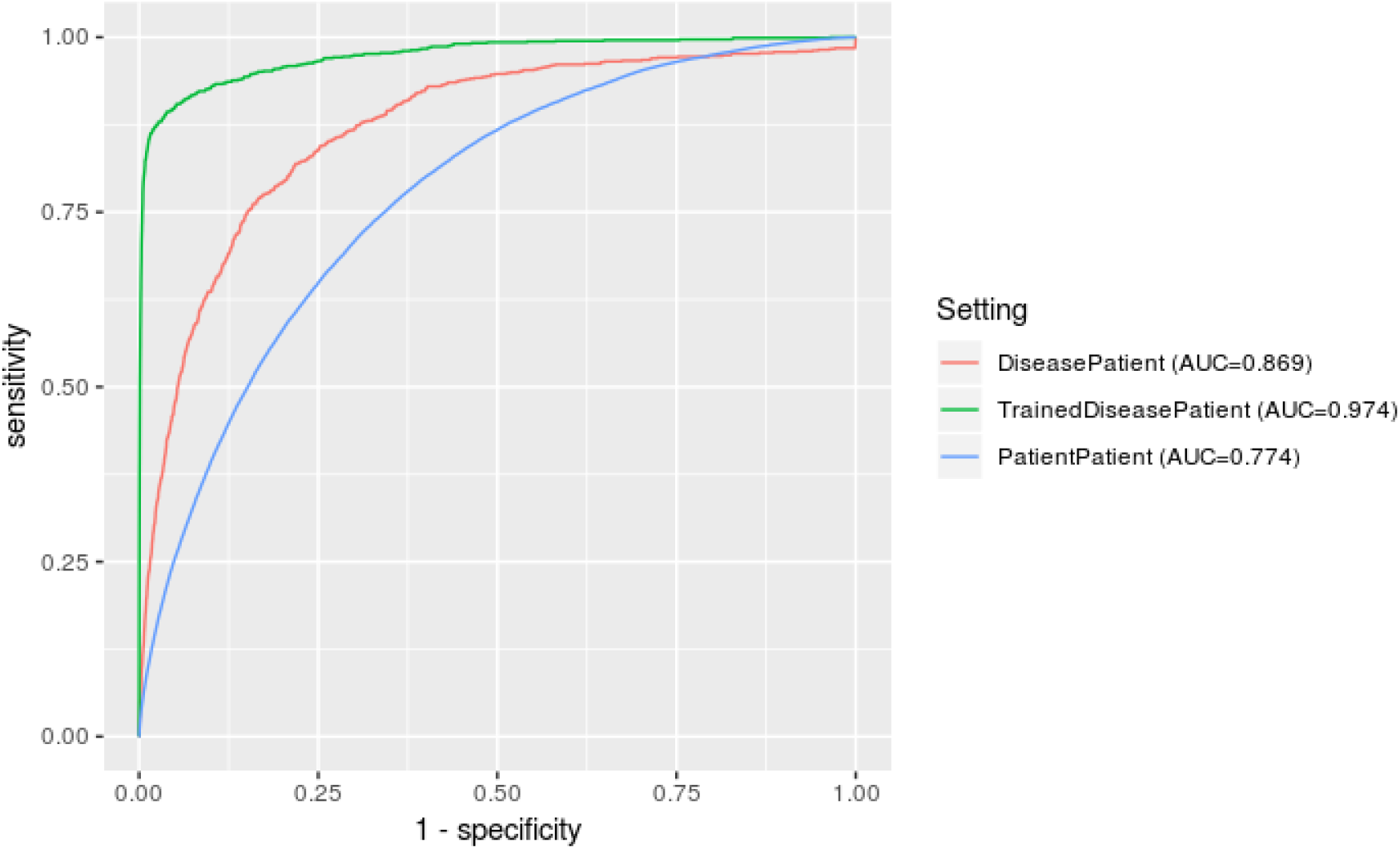
Comparative ROC curves for the models, with patient-disease comparisons showing better overall performance. The best performance was gained when adding phenotypes derived from a training set to the literature-derived phenotypes. The baseline literature disease classifier did not find all true positives, even with a false positive rate of 1, due to confidence levels of zero for certain patient-disease comparisons.

While the patient-patient comparison model yielded an inferior overall classifier, the MRR and Top Ten Accuracy metrics show that the correct disease appeared at higher ranks much more frequently than in the baseline disease definition comparison dataset. This slightly unintuitive contrast derives from two factors. The first factor consists in a difference between the number of comparisons the two models make. The patient-patient classifier compares each patient with 999 patient phenotype profiles, the disease profile classifiers compare each patient with 6,247 phenotype profiles – this provides much greater negative predictive power. That is, while the ranks of matches are higher in absolute terms for the patient-patient classifier, they are lower in relation to how many comparisons appear below it. This is compounded by the fact that patients are likely to have multiple matching patients (patients with a shared diagnosis), while there will be at most one matching disease profile.

The second factor relates to the fact that while the patient-patient model appears to have a better performance for producing highly ranked matches in absolute terms, the AUC is calculated from overall normalised semantic similarity score (becoming an overall score-derived ranking), rather than per-patient rankings (which MRR and Top Ten Accuracy are derived from). Therefore, the metric indicates that in terms of global score the patient-patient classifier is producing high scores for patient pairs that do not share a primary diagnosis. This is implied also by the difference in mean similarity score: 2.869 for the disease profile classifier, and 4.937 for the patient-patient classifier. Figure 3 also shows a comparison between the ranges of scores for the classifiers, showing that on average the patient-patient scores are both more closely clustered and greater than the patient-disease scores. This is likely due to the text annotations, and therefore the patient profiles, recording features that are irrelevant to the particular disease, but are likely to be shared across entities in the dataset. For example, given MIMIC-III is a critical care setting, many patients are suffering from or are being evaluated for pain. Across the 1,000 patient visits annotated, there are 2,380 instances of `pain (HP:0012531)`. While this would partially be controlled by the annotation frequency derived information content measure, patients who share nothing in common other than pain would nevertheless be rated more similar on that basis than those who had no annotations in common. This does not affect the disease phenotype classifier, as disease profiles have been derived from a large number of literature texts, with the final disease phenotype profiles retaining only phenotypes that appeared to be significantly associated to that disease in particular.

**Figure 3.**
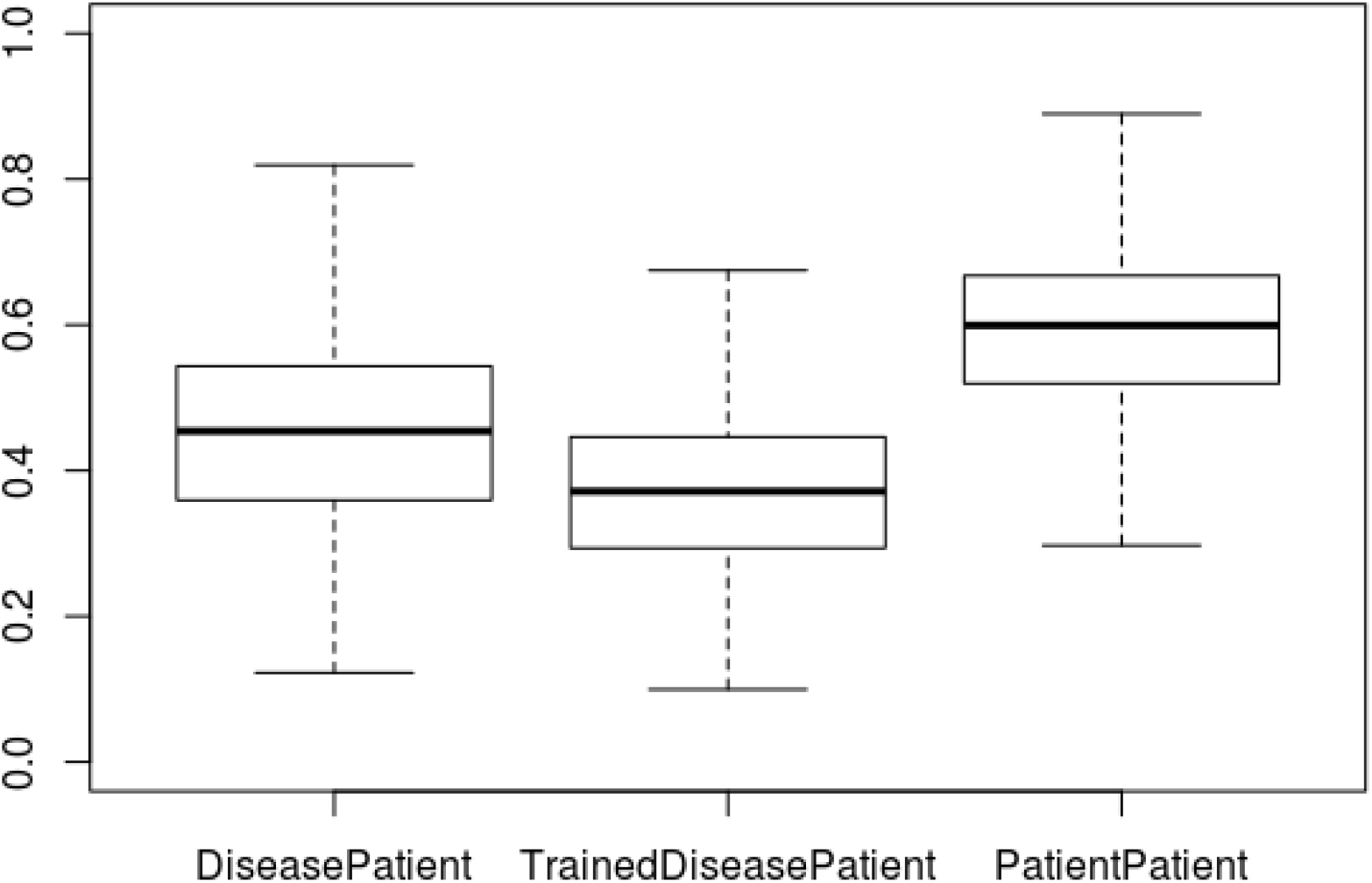
Box plot showing the different ranges of semantic similarity scores over the experiments. This shows that patient-patient comparisons were on average more similar than patient-disease comparisons, and that the extended disease profiles led to lower average semantic similarity scores than the other settings. Outliers are not shown.

The patient–patient classifier was also negatively impacted by patients who did not have another patient with a matching disease. For 107 patients, there was no true positive to be found within the ranking due to no other patient visit sharing the primary diagnosis. This situation could be improved by using a hybrid model which enhances disease phenotypes with information from patient profiles. Alternatively, a greater number of patients could be sampled, reducing the chance of an unmatched patient. To an extent, this limitation is handled within the context of the algorithm, with lower scores being assigned to patients with no matching diagnosis.

Figure 2 also shows that the baseline literature disease setting did not find all true positives. That setting, even at the highest false positive rate, was unable to recover all true positives. This is due to a proportion of patient–disease pairs being assigned with zero confidence. This happened for a matching disease profile 14 times, and in every case the disease was `‘rheumatic congestive heart failure’ (DOID:14172)`. All of the phenotypes associated with this disease, except for `sudden death HP:0001699`, were very general and therefore uninformative (with respect to the SS calculation), such as `clinical modifier (HP:0012823)` and `All (HP:0000001)`. Interestingly, this did not occur for other heart failure-related diseases, which implies that sufficiently specific and informative classes were associated with those diseases, though they did not make it to this one. This could be a potential artefact or bug in the method used to generate the literature disease phenotypes in the previous work, and could potentially be fixed by post-processing to ensure proper propagation of phenotypes across the class hierarchy.

The trained disease comparison setting, which was enhanced by phenotypes text-mined from a training set of 500 patient visits, exhibits the best performance by AUC, with the ROC curve indicating a large rise in predictive power. This indicates that the training process vastly improved the literature-derived phenotype profiles’ ability to rank patient diagnosis, bridging the qualities of the two baseline models. When compared with the patient comparison setting, this improvement comes at a small cost of precision with respect to the MRR, but with a corresponding increase in correct diagnoses appearing in the top ten. It’s possible that performance could be further improved with a larger training set, given the patient comparison relies on double the amount of patient examples. To an extent, the success of this model can be explained by the training process adapting the all diseases that appeared in the training set to the dataset – that is, all MIMIC patients will be closer to the much smaller set of diseases that appeared in the training set, and therefore to the smaller set of diseases that frequently appear in the critical care setting, even if only via phenotypes irrelevant to the disease but particular to the setting. This is evidenced by the large jump in the TPR at an extremely low FPR, visible in Figure 2. The figure also shows us that, in comparison with the baseline disease profiles, the model reaches all true positives, meaning that appropriately informative phenotypes for `‘rheumatic congestive heart failure’ (DOID:14172)` were learned through the training process.

## Discussion

We can conclude from the results that the patient-disease and patient-patient comparison models exhibit different benefits and disadvantages. The synthesis of these approaches, using patient phenotype profiles from the domain to extend literature-derived phenotypes, produced a superior model, preserving the qualities of each baseline model. This model provides strong performance in ranking primary diagnosis based on text-derived patient phenotype profiles, showing that it is a feasible and promising approach for differential diagnosis of common diseases.

Previous work in similarity-based differential diagnosis has focused either on rare diseases [8], using disease phenotype profiles taken from the OMIM database, or on manually developed phenotype profiles in particular disease domains [14, 15]. This is the first work to focus on a large set of common diseases, and therefore performance cannot be directly compared with other approaches.

Two previous works investigated the use of text-derived phenotypes for semantic similarity. Particularly, Doc2HPO created a platform for the creation of phenotype profiles, intended to be passed into other patient comparison methods. This contrasts with our approach, which used entirely uncurated text phenotypes [10]. The Doc2HPO approach also shows that curated phenotypes led to improved performance over uncurated phenotypes, although this is not a surprising result. It’s possible that future elaborations upon our work could consider different automated and semi-automated methods of optimising and curating text-derived patient phenotype profiles, outside of those provided by the semantic similarity method itself. It is also an area for future exploration to consider how the use of different text mining systems affects performance at semantic similarity tasks. Previous work comparing biomedical concept recognition tools large ranges in accuracy depending on software, approach, and parameters used [24].

The relevance of our results consist not only in the application to common diseases and text-derived phenotypes, but also in the practical secondary use of literature-derived phenotype profiles in an additional clinical setting. This extends the findings of the original work, which successfully applied the profiles for gene variant prioritisation [13]. In using those profiles for classification of diseases in this setting, we have shown that semantic similarity is amenable to transfer learning of disease phenotypes across domains.

We have also shown that out-of-domain disease profiles can be extended using examples of patient phenotype profiles derived from text in that application domain. One recent piece of work explored additions to OMIM disease phenotype profiles in the context of rare disease prediction, showing improved performance [12]. Other work has also focused on ontology extension through text, which may also improve performance of semantic similarity tasks [25]. Our previous work also showed that extension of ontologies by examining binary relations mined from text, and extension of ontology vocabularies with information from other ontologies, improved performance at a semantic similarity-based patient characterisation tasks [26, 27]. Recent work has also explored alternative methods for employing ontology axioms and taxonomy for classification and ranking problems, such as the conversion of ontology axioms to vectors [28], an approach which has been demonstrated to improve performance when compared semantic similarity approaches [29].

One limitation of our evaluation is that phenotype profiles are built from the entirety of the text record associated with the patient visit which, in most cases, will include the actual diagnosis. In that sense, it is not actually longitudinally predictive. Furthermore, while an HPO annotation refers to a phenotype, rather than a disease, there is significant cross-over between these domains, and it’s possible that the disease itself appears as a “phenotype” in the text, positively biasing the similarity task. For example, ‘hypertension’ is both the name of a phenotype and a disease, depending on the context. However, this limitation would also apply to rare diseases. Additional assessment could explore tasks in which the text record is split at the point a diagnosis was made. For example, we could identify patients who were investigated for a rare cardiac condition, splitting the text record at the point a final diagnosis was made. We could then test whether the phenotype derived from text associated with the patient prior to the final diagnosis was predictive of the true outcome.

There is also a question around the application domain for this work. Similarity-based differential diagnosis is useful for rare diseases, since they are often complex, and due to their rarity are difficult to identify by clinicians. This is less true of common diseases, especially in a critical care setting in which the diversity of diseases is limited, and are familiar to clinicians in the domain. Nevertheless, the approach still presents opportunities for clinical decision support, providing a list of conditions for a clinician to consider based on the text record. This could be particularly useful in the case of patients with rarer common diseases. The approach could also be used for automated or semi-automated coding, providing coding specialists with lists of potential codes for a given text record. The method could also be used for cohort discovery across text records, or identifying potentially misdiagnosed patients. The approach could also be extended to other outcomes, such as prognostication. For example, we could test the hypothesis that upon presentation, patients who later go on to experience more severe outcomes during a critical care visit, may be more phenotypically similar than those who do not.

The results section discussed potential difficulties in interpreting the results of the model. In previous work on similarity-based differential diagnosis for rare genetic diseases, p-values were associated with similarity scores, providing a clearer way of interpreting results [8]. It is also possible that front-end user-facing software, such as Phenomizer [8], could be re-used for this purpose, extending it with a larger set of phenotypes and disease profiles. This could help to move our approaches towards practically useful clinical tools.

In terms of improving performance and exploring different settings, we could explore different methods of calculating similarity or information content. For example, the current measure of information content is weighted by the frequency the concept appears in the corpus, however these can also be calculated topologically, on the basis of how general or specific the classes are [30]. We consider it an area for future work to create an easily modifiable platform for comparison of different settings in this problem domain, with associated benchmarks.

## Conclusions

We explored methods for similarity-based classification and differential diagnosis of patient critical care visits using text-derived phenotypes. In baseline experiments, comparing patient phenotype profiles with profiles derived from literature produced a better classifier, though comparing patients with other patients was far more likely to produce highly ranked matches. We then combined these two methods, extending literature-derived phenotype profiles with patient phenotypes mined from text associated with MIMIC patient visits from a training set. This produced the superior classifier, retaining the negative predictive power of the literature-derived phenotypes, producing also the highly ranked matches of the patient-patient comparisons. We believe this is a promising development, and could lead to powerful semantic similarity derived solutions for differential diagnosis of common diseases or outcomes, as well as text classification and cohort discovery tasks in a clinical context.

## Ethical approval

This work makes use of the MIMIC-III dataset, which was approved for construction, de-identification, and sharing by the BIDMC and MIT institutional review boards (IRBs). Further details on MIMIC-III ethics are available from its original publication (DOI:10.1038/sdata.2016.35). Work was undertaken in accordance with the MIMIC-III guidelines.

## Availability of data and material

Komenti is an open source text mining framework, and is available under an open source licence from https://github.com/reality/komenti. The code used to sample MIMIC, and run similarity experiments is available from https://github.com/reality/miesim/. The annotated MIMIC dataset is not made publicly available, because researchers are required to meet ethical conditions to access MIMIC-derived datasets. To access this dataset, please contact the corresponding author directly.

## Acknowledgements

GVG and LTS acknowledge support from support from the NIHR Birmingham ECMC, NIHR Birmingham SRMRC, Nanocommons H2020-EU (731032) and the NIHR Birmingham Biomedical Research Centre and the MRC HDR UK (HDRUK/CFC/01), an initiative funded by UK Research and Innovation, Department of Health and Social Care (England) and the devolved administrations, and leading medical research charities. The views expressed in this publication are those of the authors and not necessarily those of the NHS, the National Institute for Health Research, the Medical Research Council or the Department of Health. RH and GVG were supported by funding from King Abdullah University of Science and Technology (KAUST) Office of Sponsored Research (OSR) under Award No. URF/1/3790-01-01. AK was supported by by the Medical Research Council (MR/S003991/1) and the MRC HDR UK (HDRUK/CFC/01).

